# GANai: Standardizing CT Images using Generative Adversarial Network with Alternative Improvement

**DOI:** 10.1101/460188

**Authors:** Gongbo Liang, Sajjad Fouladvand, Jie Zhang, Michael A. Brooks, Nathan Jacobs, Jin Chen

## Abstract

Computed tomography (CT) is a widely-used diag-reproducibility regarding radiomic features, such as intensity, nostic image modality routinely used for assessing anatomical tissue characteristics. However, non-standardized imaging protocols are commonplace, which poses a fundamental challenge in large-scale cross-center CT image analysis. One approach to address the problem is to standardize CT images using generative adversarial network models (GAN). GAN learns the data distribution of training images and generate synthesized images under the same distribution. However, existing GAN models are not directly applicable to this task mainly due to the lack of constraints on the mode of data to generate. Furthermore, they treat every image equally, but in real applications, some images are more difficult to standardize than the others. All these may lead to the lack-of-detail problem in CT image synthesis. We present a new GAN model called GANai to mitigate the differences in radiomic features across CT images captured using non-standard imaging protocols. Given source images, GANai composes new images by specifying a high-level goal that the image features of the synthesized images should be similar to those of the standard images. GANai introduces an alternative improvement training strategy to alternatively and steadily improve model performance. The new training strategy enables a series of technical improvements, including phase-specific loss functions, phase-specific training data, and the adoption of ensemble learning, leading to better model performance. The experimental results show that GANai is significantly better than the existing state-of-the-art image synthesis algorithms on CT image standardization. Also, it significantly improves the efficiency and stability of GAN model training.

## I. Introduction

Computed tomography (CT) is one of the most popular diagnostic image modalities routinely used for assessing anatomical tissue characteristics for disease management [1], [2], [3], [4], [5]. CT scanners provide the flexibility of customizing acquisition and image reconstruction protocols to meet an individual’s clinical needs [6], [7]. However, CT acquisition parameter customization is a double-edged sword [8]. While it enables physicians to capture critical image features towards personalized healthcare, it forms a barrier to analyzing CT images in a large scale, in that capturing CT images with non-standardized imaging protocols may result in inconsistent radiomic features [9], [10]. As was revealed in a recent study, both intra-CT (by changing CT acquisition parameters) and inter-CT (by comparing different scanners with the same acquisition parameters) tests have demonstrated low shape, and texture, for CT imaging [11], [12]. In the example shown in Figure 1, each lung tumor was acquired twice using two different reconstruction kernels (Bl64 and Br40, Siemens Healthineers, Erlangen, Germany). The figure demonstrates that the appearances (as well as the radiomic features) of the same tumor can be strongly affected by the selection of CT acquisition parameters.

**Fig. 1:**
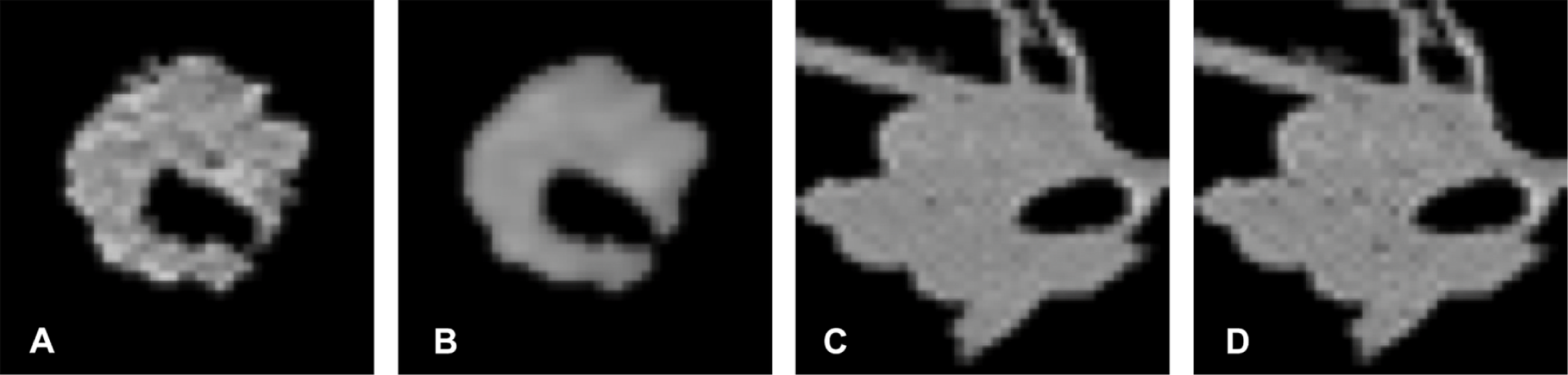
Lung tumors acquired using two kernels have shown significantly different appearances as well as radiomic features. (A) Lung tumor 1 acquired with kernel Bl64. (B) Lung tumor 1 acquired with kernel Br40. (C) Lung tumor 2 acquired with kernel Bl64. (B) Lung tumor 2 acquired with kernel Br40.

To overcome the barriers that prevent the use of CT images in large-scale radiomic studies, algorithms have been developed aiming to integrate and standardize CT images from multiple sources. Image synthesis is a class of algorithms that generate synthesized images from source images, which satisfy the condition that the feature-based distributions of the synthesized images are similar to that of target images [13]. Mathematically, given source image *x*, an image synthesis algorithm composes a synthesized image *x*′ by specifying a *high-level* goal that the image features of *x*′ are significantly more similar to that of the target image *y* than the source image *x*. Image synthesis algorithms have been widely used in image conversion and natural language processing, such as the synthesis of images from text descriptions [14]. Note that image synthesis is different from image conversion (such as to convert an MRI image to a CT image), which requests an exact pixel-to-pixel match between the synthesized images and the target images [15].

Image synthesis algorithms can be roughly classified into two groups, i.e., traditional image processing algorithms and deep learning-based algorithms. In the first group, the histogram matching-based algorithm has been widely used [16], [17], [18], [19]. In general, it synthesizes images by mapping the histogram of source images to that of target images. However, finding the mapping function requires the presence of the target images, which are often missing or are not well defined in practice. In the second group, generative adversarial network models (GAN), a class of deep learning algorithms, can learn the data distribution of training data and generate synthesized examples which fall under the same distribution of the training [20]. In particular, the conditional generative adversarial network (cGAN), a special kind of GANs, learns the conditional distribution of the source image *x* given the target image *y* and then performs image transference from one domain to another [21], [22]. However, GAN models (include cGAN) are not directly applicable to our task mainly due to three limitations: 1) GAN models do not contain any constraints to control what modes of data it shall generate; 2) the synthesized images are not guaranteed to be similar to the target images (Figure S1); 3) GAN models treat every image in training equally, but in real applications, some images are more difficult to synthesize than the others (Figure S2). All these limit the functionality of GAN models and may lead to the lack-of-detail problem in image synthesis.

To address the computational challenges in medical image synthesis, where great image details have to be maintained, we propose a novel deep learning framework called “Generative Adversarial Network with Alternative Improvement (GANai)”. GANai has a similar architecture as cGAN, but its training process is significantly different. Specifically, GANai introduces an alternative improvement training strategy to alternatively train its deep learning components and steadily improve the whole model performance. The adoption of the new training strategy enables a series of technical improvements, including phase-specific loss functions, component-dedicate training data, adoption of ensemble learning, and so on, leading to a significant improvement on model performance.

While GANai can be deployed in many applications, we adopted and evaluated GANai in mitigating the differences in radiomic features due to using non-standardized CT imaging protocols. The experimental results show that GANai is significantly better than the state-of-the-art image synthesis algorithms, such as cGAN and histogram matching, on all the image acquisition parameters that we have tested. In summary, GANai has the following computational advantages:

1. GANai introduces an alternative improvement training strategy to alternatively and steadily improve model performance.
2. GANai adopts a new phase-specific loss function that allows the discriminator and the generator to collaborate rather than competing with each other.
3. GANai improves model training effectiveness by training the discriminator and the generator using specified training images.
4. GANai adopts ensemble learning to significantly improve the stability of GAN model training.

## II. Background

Radiomics is an emerging science to extract and use comprehensive radiomic features from a large volume of medical images for the quantification of overall tumor spatial complexity and the identification of tumor subregions that drive disease transformation, progression, and drug resistance [23], [24], [25], [26]. However, due to the use of non-standardized imaging protocols, variations in acquisition and image reconstruction parameters may cause inconsistency in radiomic features extracted from images, which poses a barrier to the practice of radiomics in large-scale [10], [24], [25].

### A. CT Image Acquisition Parameters

In modern CT imaging, there are a large number of imaging protocols, and using non-standardized imaging protocols is common [6]. The CT image acquisition parameters includ kV (the x-tube voltage), mAs (the product of x-ray tube current and exposure time), collimation, pitch, reconstruction kernel, field-of-view, and slice thickness [27], [28]. In routine clinical practice, certain parameters are often adjusted to meet the diagnostic needs, i.e., to obtain satisfactory image quality while maintaining low radiation dose to patients. Changing acquisition parameters may significantly affect the resulting images (Figure 1). For example, adjusting kV will change CT numbers (the pixel values of a CT image), changing mAs will affect image noise rate, and the selection of reconstruction algorithms will result in different image texture features.

### B. Histogram Matching

Histogram matching (or called histogram specification) is a widely-used image synthesis tool. It uses the intensity histogram to represent images and then transforms a source image to a target image by matching their intensity histograms [16], [17], [18], [19]. While histograms can represent the density of intensity in the whole image, the major drawback is the loss of location information. A variation of histogram matching is to divide a source image into multiple patches and to apply histogram matching on each patch, expecting that such patch-based representation may lead to location-specific image synthesis. However, patch-based histogram matching may introduce artifacts, esp. on the edges of patches. It is also sensitive to the selection of matching parameters such as the number of bins of a histogram (Figure S3).

### C. Generative Adversarial Networks

Recently, deep learning has shown remarkable performance in various medical informatics tasks. For example, it has surpassed the human experts’ performance on skin cancer classification by only looking at the dermoscopic images [29].

The generative adversarial network (GAN) is a kind of deep learning models that learns the data distribution of training images and generate synthesized images under the same distribution [20], [30], [31]. A GAN model usually has two components, i.e., the discriminator *(D*) and the generator (G), where *G* generates synthesized data from random noise, and D learns a data distribution from the training data and determines whether the synthesized data generated by G fall into the distribution. The goal of G is to generate synthesized data which are good enough to fool D, while D always aims to discriminate the synthesized data and the real data.

The conditional generative adversarial network (cGAN) is a kind of GAN models that learns the *conditional* distribution of the training data and generates synthesized data under the same condition [21], [32], [33]. Among cGAN models, the Image-to-Image model performs the image-to-image transference from one domain to another concerning the given condition, and it has become a widely recognized conditional image synthesis model [22]. Note that the images synthesized by cGANs are not necessarily similar to the target images, although they look “real”, meaning having similar semantic meanings as the target images (see Figure S1, S2). However, in medical applications, it is important to maintain authenticity in the synthesized CT images. Specifically, it is expected to generate images with the distribution of radiomic features significantly similar to that of the target images.

While GAN models are advanced in image synthesis [22], [14], image inpainting [34], semantic segmentation [35], etc., GAN models are suboptimal regarding training efficiency and stability. To address the GAN training problem, several ensemble learning-based strategies have been applied to improve model training: 1) to train multiple GANs in parallel using a random initialization of model parameters, and then to randomly choose one of the GANs to generate the synthesized data [36]; 2) to train multiple Ds and requires the G to fool a group of Ds [37]; and 3) to select training data using boosting and to train a cascade of GANs in sequence. It has been shown that the performance of GANs can be significantly improved by using ensemble learning [37].

## III. Method

To extend the adversarial learning into the medical image domain and to address the aforementioned challenges, we propose Generative Adversarial Network with Alternative Improvement (GANai).

### A. Architecture

GANai consists of two components, i.e., the generator (*G*) and the discriminator (*D*), where *G* is a U-Net with fifteen hidden layers and *D* is a multilayer perceptron model with six fully connected layers [38]. The architecture of GANai is similar to the cGAN models, shown in Figure 2 [22]. The inputs of *D* of GANai are image pairs (*x*, *y*) and (*x*, *x*′), where (*x*, *y*) denotes the real pair (positive training), and (*x*, *x*′) denotes the fake pair (negative training). The goal of *D* is to distinguish the real pairs from the fake pairs. Given the feedback from *D*, *G* learns the mapping from *X* to *Y* and generates a synthesized image *x*′ for any given source image *x*(*x* ∈ *X*) in *Y*’s domain. In contrast to *D*, *G* aims to synthesize images that can fool *D*. If *D* can distinguish most of the fake pairs from the real pairs, the performance of *G* needs to be further improved. Otherwise, we conclude that the generative results of *G* are good enough for the current *D*.

**Fig. 2:**
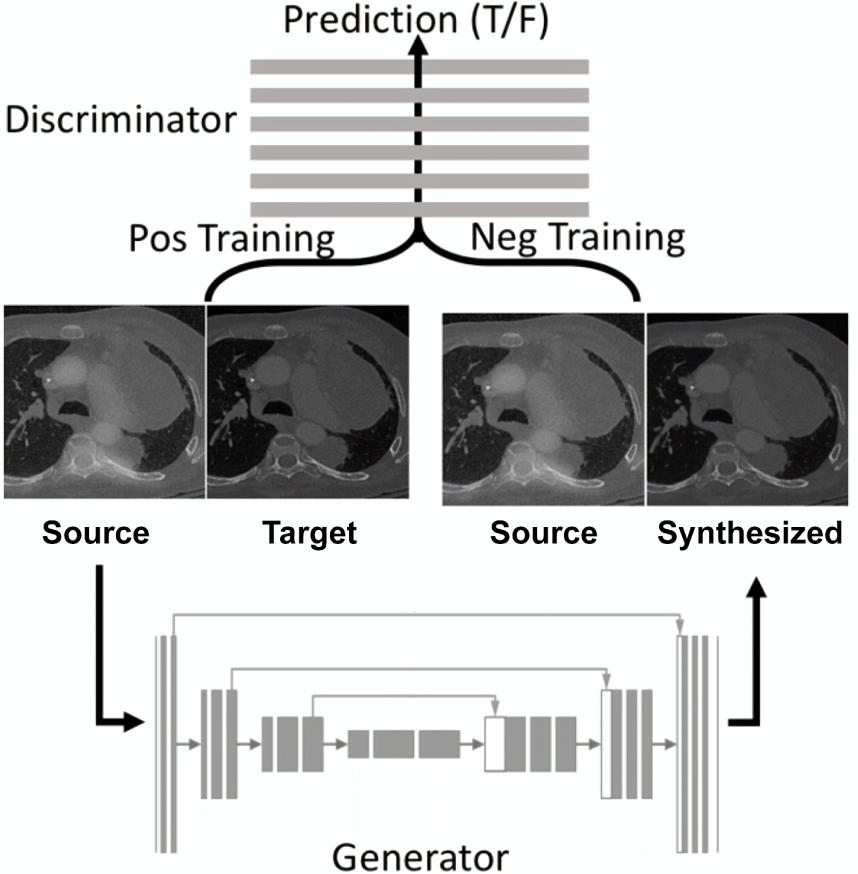
Architecture of GANai. Given a source image, the generator *G* synthesizes a new image to fool the discriminator *D*, while *D* aims to distinguish the synthesized image and the target image.

### B. Alternative Improvement

In traditional GAN models, *D* and *G* are trained synchronously (*D* and *G* trained together) or asynchronously (several batches of *D*-training followed by several batches of *G*-training), based on the assumption that both *D* and *G* can be gradually improved together. In practice, however, if *D* is not well trained to capture the intrinsic features to separate a real and a fake image, *G* can easily fool *D*. Similarly, if *G* is not well “challenged” by *D*, its model performance is not guaranteed to be improved.

We introduce the alternative training approach for GANs (Figure 3). As the name suggested, GANai has two alternate training phases, i.e., the discriminator training (*D*-training) and the generator training (*G*-training). In each training phase, we focus on optimizing one of the components while freezing the other. A training phase will stop if the current component is well trained or the training step exceeds an upper bound (see Section V-B for more details). After that, we switch to the other training phase (Figure 3 solid lines). The alternative training strategy enables a series of technical improvements, including phase-specific loss functions, phase-specific training data, and the adoption of ensemble learning, which will be introduced in the following subsections.

**Fig. 3:**
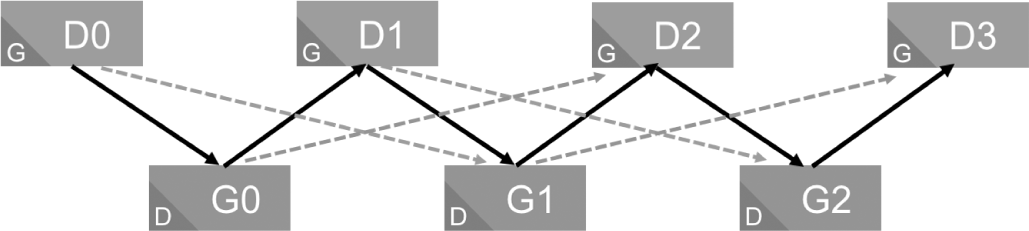
In each training phase of GANai, D (or *G*) is trained while the other component is frozen. The name of a block (such as Do or Gi) indicates the component in training, and the letter located at the bottom left corner indicates the component that is frozen. The alternative training (solid line) ensures high performance while the ensemble approach (dotted line) improves the training stability.

### C. Loss Functions

The alternative training of GANai may boost model performance by preventing each component being too strong or too weak. In the literature, strategies have been presented to freeze part of a GAN when the GAN components are imbalanced [39]. However, it is difficult to decide when to freeze/unfreeze a component of GAN. To address this issue, we redesigned the loss functions.

In the *D*-training phase, *G* is frozen so that *D* learns the differences between the synthesized images and the target images and discriminates the synthesized images. Hence, the loss function of *D* is the same discriminator loss of cGAN [21]:

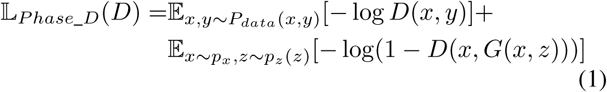

where *x* is the source image; *y* is the target image; *G*(*x*, *z*) is the synthesized image generated by *G*, which maps the source image *x* and a random noise vector *z* to *y*; *D*(*x*, *y*) is the prediction result of the real pair; and *D*(*x*, *G*(*x*, *z*)) is the prediction result of the fake pair. For *D*(*x*, *y*), the higher the prediction accuracy, the higher the value of *D*(*x*, *y*).

In the *G*-training phase, *D* is frozen, and it evaluates the results of *G*. Since we expect *G* to fool *D*, the loss of *D* in the *G*-training phase is defined as:

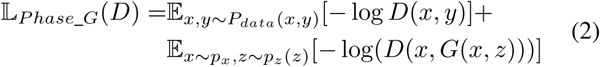

Finally, by integrating Eq 1 and Eq 2, the loss function of *D* in GANai is defined as:

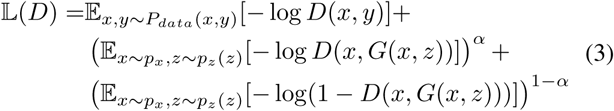

where parameter α = 1 if GANai is in the *G*-training phase and α = 0 in the *D*-training phase.

The loss function of *G* is the same as Isola et al. [22]. Also, we adopt the *L*1 loss as the regularization factor.

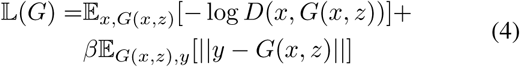

where *β* is the weight of the regularization term.

To determine when to switch between the *D*-training phase and the *G*-training phase, the prediction accuracy on the fake image pairs (*D*(*x*, *x*′)) is used. The value of *D*(*x*, *x*′) is computed at every training step and is compared with two thresholds. More specifically, if *D*(*x*, *x*′) < *T*_l_, GANai will switch from *D*-training to *G*-training. If *D*(*x*, *x*′) > *T*_*h*_, GANai will switch from *G*-training to *D*-training. *T*_*l*_ and *T*_*h*_ are the lower and upper thresholds of *D*(*x*, *x*′). To improve training stability, the least amount of steps (minibatches) of each training phase is also specified. Note that in GANai, the value of *D*(*x*, *x*′) increases and decreases, indicating that the performance of *D* and *G* is improved alternatively.

### D. Training D and G with Dedicated Training Data

Since the components of GANai are trained separately, one idea is to increase model training efficiency by training *G* and *D* using different data. More specifically, the images that are potentially synthesizable can be used to accelerate the *G*-training, while the training of *D* can benefit from images that are difficult to synthesize.

We develop a procedure to select training data for *D* and *G*. First, a cGAN model is trained using all the training data [22]. Second, with the trained cGAN model, we synthesize a new image for every source image and compare every synthesized image with its corresponding target image using Kullback-Leibler divergence [40], normalized mutual information (NMI) [41], and cosine similarity. Finally, the training data is split into two subsets based on z-score, i.e., 1/3 of the source-target image pairs with the highest similarities between synthesized images and target images (called *T*_*easy*_) and 1/3 of the images with the lowest similarities (call *T*_*hard*_). The new procedure allows us to train G using *T*_*easy*_ and train *D* using *T*_*hard*_ (see Section V-A for other training set selection strategies).

### E. Improving Training Stability using Ensemble Learning

Due to the nature of the generative adversarial concept (i.e., open-ended competition between GAN components), it is not guaranteed that *G* or *D* will improve towards the same direction. For example, if the kth state of *G* fools the (*k* − 1)th state of *D*, it still may be classified by the older (*k* − 2)th state of *D*. Therefore, during the two-phase training of GANai, we improve the model stability by adopting the ensemble learning. Simply speaking, a *D* is required to discriminate multiple *G*s and a *G* must fool multiple *D*s.

Mathematically, the following criteria are specified in GANai: when training the *k*th *G*, the *G* must fool both the (*k* − 2)th state and the (*k* − 1)th state of *D*, and when training *k*th *D*, the *D* should discriminate both the (*k* − 2)th state and the (*k* − 1)th state of *G*. For an illustrative example, see the dot lines in Figure 3. These criteria can be further extended to incorporate more historical *D*s or Gs or more sophisticated conditions. In the exception that GANai cannot identify such a *D* or *G* that satisfies the criteria after at most *T*_*s*_ steps (the maximum training step in each phase), it will roll back to the previous state, and re-train the current component.

## IV. Experimental Results

### A. Data

In total 2,448 chest CT image slices of lung cancer patients were collected using Siemens CT Somatom Force at the University of Kentucky Medical Center. For each patient, a CT image was constructed with each of the possible combinations of two image reconstruction parameters, i.e., slice thickness (0.5, 1, 1.5, 3mm) and reconstruction kernels (Bl57 and Bl64). With data augmentation, the training data has been extended to 14,958 image patch pairs. Among them, 7,479 assigned as *T*_*easy*_ and 7,479 assigned as *T*_*hard*_ using the procedure introduced in Section III-D. Each image pair contains a source image *x* and the target image y. See details of data augmentation in Section S1.A with examples in Figure S4.

The validation data contains 3,554 2.5D images, and multiple radiomic features were extracted for model validation. Specifically, we randomly cropped 2.5D images from the CT images that have not been used as training data, with their dimensions ranging from 5 × 5 × 5 to 60 × 60 × 30 pixels. When cropping the 2.5D validation images, we excluded areas with bone or air, since soft tissues are what physicians are most interested. See Section S1.B for more details.

Given a large number of CT imaging protocols, it is impractical to apply all of them. We selected two image reconstruction parameters (kernel and slice thickness) and used all the combinations for the model performance test. Also, we chose 1mm slice thickness and Bl64 kernel to be the standard imaging protocol, since it is widely used in the current lung cancer radiomic studies. Note the settings can be easily extended to incorporate more acquisition parameters or to use a different standardized imaging protocol.

### B. Implementation Details of GANai

In GANai, *G* is a fifteen hidden layers U-Net [38], with the size between 128 × 128 × 64 and 1 × 1 × 512 (Figure S5). The input of G are 256 × 256 images, and the synthesized images have the same image size. *D* is implemented as a multilayer perceptron model with six fully connected layers with the size between 256 × 256 × 3 and 30 × 30 × 1 (Figure S6).

The training of GANai started with the *D*-training phase, and all the network weights were randomly initialized. We set the regularization term weight β = 100 to reduce the visual artifacts [22], and used *T*_*l*_ = 0.05 and *T*_*h*_ = 0.95 as the training phase switch thresholds, and *T*_*s*_ = 10 epochs as the maximum training step. Within each training phase, the model needed to be trained for at least five steps before switching to the other training phase. GANai was trained for 100 epochs with learning rate being 0.0002, momentum being 0.5.

GANai is deployed on Tensorflow [42] on a Linux computer server with eight Nvidia GTX 1080 GPU cards. It took 15 hours to train GANai from scratch using a single GPU card. Using the trained model, it took 0.2 seconds to generate a synthesized image (5 images per second).

Figure 4 shows the discriminator prediction results on all the fake pairs *D*(*x*, *x*′) in the first 150 steps of training. With the training of *D*, *D*(*x*, *x*′) decreases. When the value of *D*(*x*, *x*′) is below *T*_*l*_ (in our experiment, *T*_*l*_ = 0.05), GANai is switched to the *G*-training phase. In the *G*-training phase, *D*(*x*, *x*′) increases, since *D* is frozen and the performance of *G* keeps increasing. When the value of *D*(*x*, *x*′) is higher than *T*_*h*_ (*T*_*h*_ = 0.95), GANai is switched to the *D*-training phase.

**Fig. 4:**
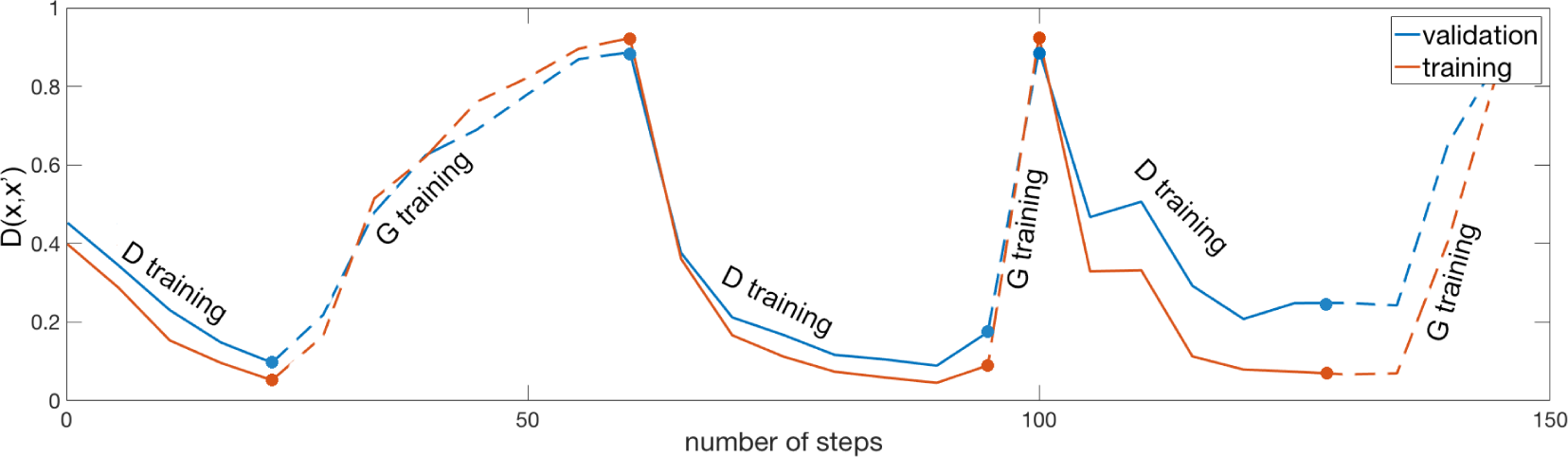
The prediction results of *D* on the fake image pairs (*x*, *x*′) in the first 150 steps of the alternative training. For *D*(*x*, *x*′), the higher the prediction accuracy, the lower the value (*D*(*x*, *x*′) ∈ [0, 1]).

The training and validation loss of *D* and *G* in the first 150 training steps are shown in Figure 5. Both the training and validation loss of *D* decreased in every training phase, which indicates the model performance of *D* and *G* was improved alternatively. In the *D*-training phase, if the performance of *D* is increased, the loss of *D* will reduce, since both −log(*D*(*x*, *y*)) and − log(1 − *D*(*x*, *x*′)) are both reduced (solid lines in Figure 5A). When switching from the *D*-training to the G-training phase, α in the loss function of *D* flips from 0 to 1, which immediately turns the loss of *D* from a small value to a high value (see the jumps located at phase turning points in Figure 5A). In the *G*-training phase, if the performance of *G* is increased, the performance of *D* will decrease, so the loss of *D* decreases (dotted lines in Figure 5A). Figure 5B shows the loss of *G* increases in *D*-training phase (due to the performance improvement of *D*) and decreases in *G*-training phase, since the performance of *G* is improved (See Section V-C).

**Fig. 5:**
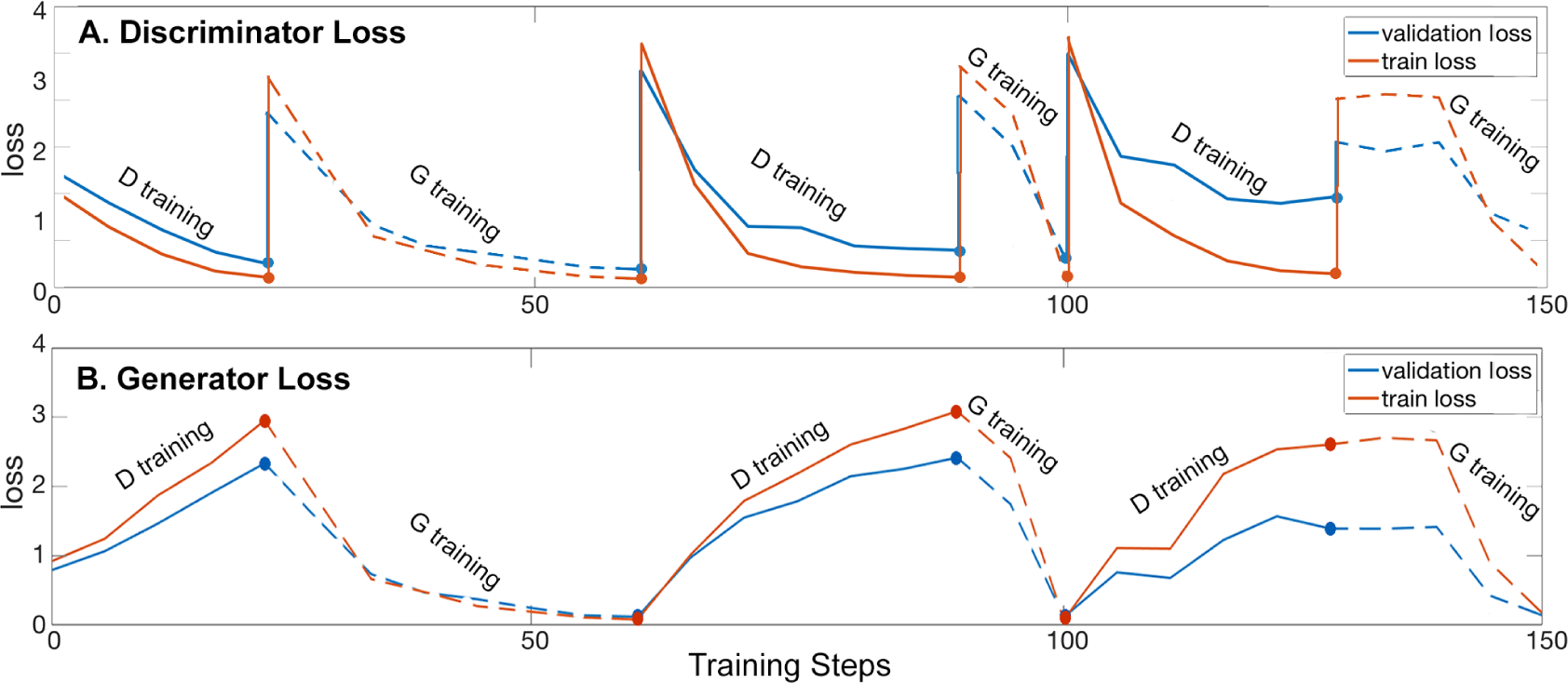
The training loss and the validation loss of *D* and *G* in GANai in first 150 steps of training. The solid lines indicate the loss of *D* in the *D*-training phase. The dotted lines indicate the loss of *D* in the *G*-training phase. The solid points indicate the time when GANai switches between the *D*-training phase and the *G*-training phase.

### C. Evaluation Metric

For performance evaluation, we compared GANai with cGAN [22] and the patch-based histogram matching (see details in supplementary section III). Instead of hiring human annotators, we adopt the radiomic features for performance evaluation [43], [44]. Specifically, two classes of radiomic features were used for model performance evaluation, i.e., 2.5D texture features (i.e., gray-level co-occurrence matrix) and 2.5D intensity histogram based features. In total, eight radiomic features were adopted for performance evaluation (see Section S2 for details).

Per every radiomic feature to test, we compared each synthesized image and its target image, and computed the absolute error and relative error using the following equations:

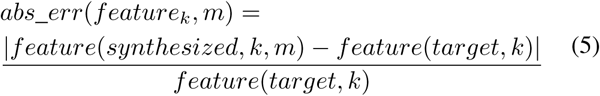

where *feature*_*k*_ is the *k*th radiomic feature, *m* is either GANai or a image synthesis model to compare.

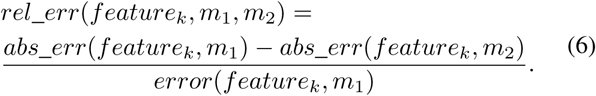

where *m*_1_ and *m*_2_ are two different image synthesis models. For the relative error, a positive value indicates that *m*_2_ has smaller error than *m*_1_, vice versa.

Model stability is evaluated using the cumulative sum control chart (CUSUM) [45]. CUSUM is a sequential analysis model typically used for monitoring change detection [46]. In CUSUM, the differences between any two adjacent values (in our case, the absolute errors between any two adjacent saved model states) are measured and are compared with a threshold. CUSUM is computed as the number of the difference values higher than a threshold (called out-of-control points). In our experiment, a series of CUSUM values were generated for each model using multiple thresholds. The normalized sum of the CUSUM values, which is the smaller the better, was used for model stability evaluation.

### D. Performance Evaluation Results on Generator

The absolute errors on all the tested radiomic features are shown in Table I. For the detailed feature-based errors, see Figure S7. On the texture features, the mean absolute error of histogram matching over all six features is 0.37. cGAN reduces it to 0.13, and GANai further reduces the absolute error significantly to 0.08 (two sample t-test *p* − *value* ≤ 0.01). On the intensity histogram features, GANai decreases the absolute errors by 17.77% from cGAN, and 79.05% from histogram matching. The results indicate that GANai is significantly better than cGAN and patch-based histogram matching.

**TABLE I:** Averaged absolute errors (SD) of (1) the texture features and (2) the intensity histogram features computed using histogram matching, cGAN, and GANai. In all of them, GANai has the smallest errors (cGAN and GANai two sample t-test *p* − *value* ≤ 0.01).

| Absolute Error | Hist. Matching | cGAN | GANai |
| --- | --- | --- | --- |
| Contrast<sup>1</sup> | 0.21 ± 0.15 | 0.12 ± 0.08 | **0.09** ± 0.06 |
| Correlation<sup>1</sup> | 0.18 ± 0.13 | 0.18 ± 0.12 | **0.09** ± 0.07 |
| Dissimilarity<sup>1</sup> | 0.15 ± 0.11 | 0.09 ± 0.06 | **0.06** ± 0.04 |
| Energy<sup>1</sup> | 0.47 ± 0.28 | 0.19 ± 0.14 | **0.14** ± 0.11 |
| Entropy<sup>1</sup> | 0.09 ± 0.06 | 0.02 ± 0.01 | **0.01** ± 0.01 |
| Homogeneity<sup>1</sup> | 0.28 ± 0.16 | 0.10 ± 0.06 | **0.07** ± 0.05 |
| Kurtosis<sup>2</sup> | 0.54 ± 0.27 | 0.18 ± 0.14 | **0.15** ± 0.11 |
| Skewness<sup>2</sup> | 0.51 ± 0.27 | 0.16 ± 0.12 | **0.14** ± 0.11 |

Table II shows the relative errors of GANai and cGAN on seven sets of the validation data generated using different combinations of CT acquisition parameters. A positive value indicates the error of GANai is lower than cGAN, while a negative value indicates the error of GANai is higher than cGAN. The results show that GANai outperforms cGAN on five out of seven validation subsets, on which GANai decreased the relative errors by 36.21% on average. For example, on the texture features, GANai reduces the relative error by 54.48% on the Bl64 kernel with 0.5mm slice thickness images. For the detailed feature-based errors, see Figure S8-S15.

**TABLE II:**
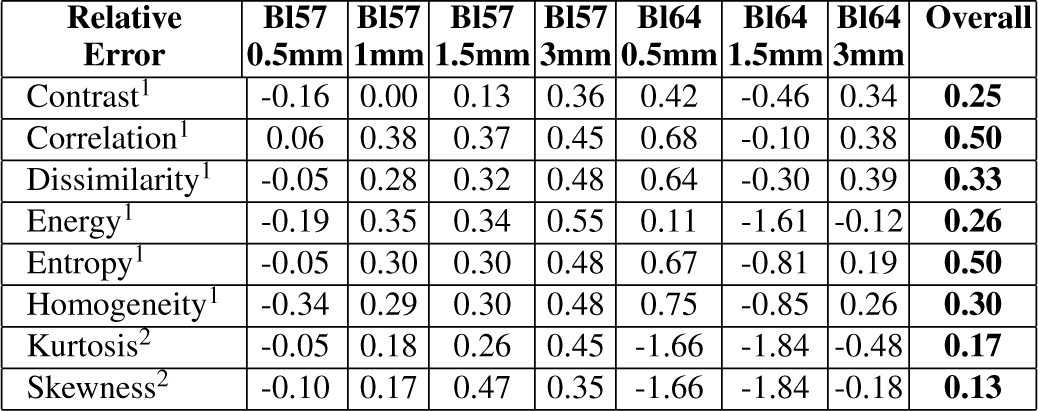
Averaged relative errors on the texture features^1^ and the intensity histogram features^2^ by comparing cGAN and GANai. Positive values mean GANai is better, and negative values mean cGAN is better. Overall, GANai has smaller errors than cGAN.

Figure 6 shows an example of the synthesized images using cGAN or GANai generated after 100 training epochs. The GANai synthesized image is more similar to the target image, has sharper edges, and has fewer artifacts than cGAN. Figure 7 shows both cGAN and GANai model reaches their best performance after 20 epochs of training. After that, GANai can still maintain high synthesized image quality, but cGAN started to introduce artifacts.

**Fig. 7:**
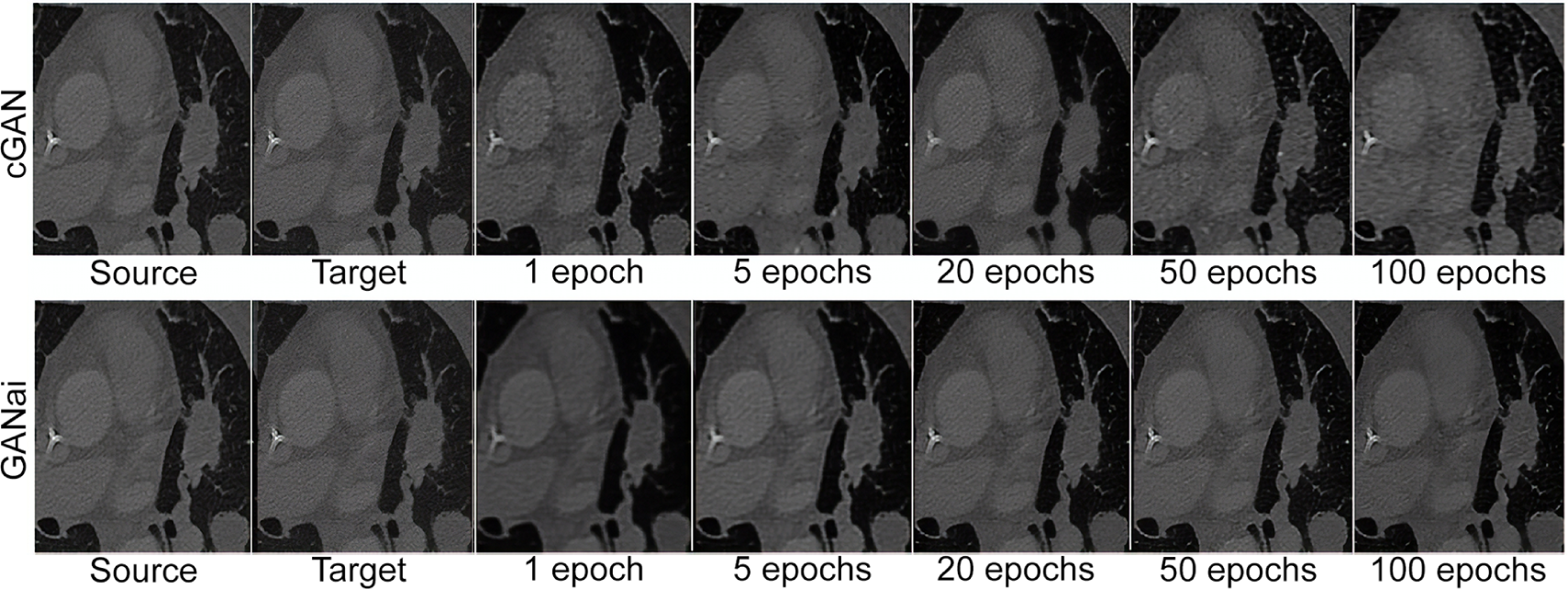
Examples of the synthesized images generated by cGAN and GANai at multiple training steps. The first two columns are the source and target images. Both cGAN and GANai reached their best performance at about 20 training epochs. The synthesized images generated by cGAN have obvious artifacts and have less sharp edges than that of GANai. Furthermore, GANai maintained a high synthesized image quality in the continuous training after the first 20 epochs, whereas cGAN started to introduce additional artifacts into the synthesized images.

### E. Performance Evaluation Results on Discriminator

To evaluate the performance on the discriminator *D*, we generated a fake-pair-only dataset and used it to measure the prediction accuracy of all the *D*s in the model training process. Specifically, given a fixed source image set *X*_*val*_ and the correspondent target image set *Y*_*val*_, each having 1,750 images, we generated the synthesized image set 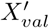 using the second last generator of GANai. The accuracy of every discriminator (such as *D*_0_ to *D*_3_ in Figure 3) in the alternative training process of GANai was measured with all image pairs in 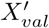. Accuracy is defined as the proportion of (*x*, *x*′) that were correctly classified as the fake image pairs. Figure 8 shows the prediction accuracy of *D* at every training process. The increasing prediction accuracy shows the performance of *D* was steadily improving during the training of GANai.

**Fig. 8:**
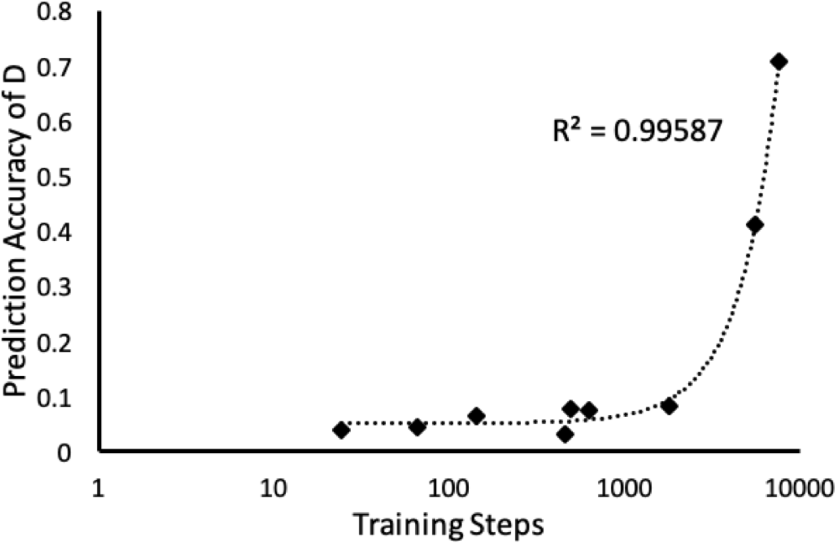
Prediction accuracy of *Ds* gradually increases during the alternative training process of GANai.

### F. Performance Evaluation Results on Training Stability

In GANai, an ensemble learning-based approach is adopted to increase the training stability. To demonstrate the effectiveness of this approach, we designed the following experiment. Three networks (cGAN, GANai*ai*_*singleDG*_, and GANai) were trained for 100 epochs using the same training data, where *GANai*_*singleDG*_ is a simplified version of GANai that trains the current component only based on the previous counter component, without using multiple *D*s or *G*s. The training state of every 2.5 training epochs was saved. We compared all the three models using the same validation data at every saved model state (Figure 9A). The normalized sum of the CUSUM values of cGAN, *GANai*_*singleDG*_, and GANai over all the six texture features are 0.21, 0.15, and 0.13 respectively, indicating GANai is the most stable model among the three. Figure 9 shows the CUSUM on the contrast feature computed using the gray-level co-occurrence matrix.

**Fig. 6:**
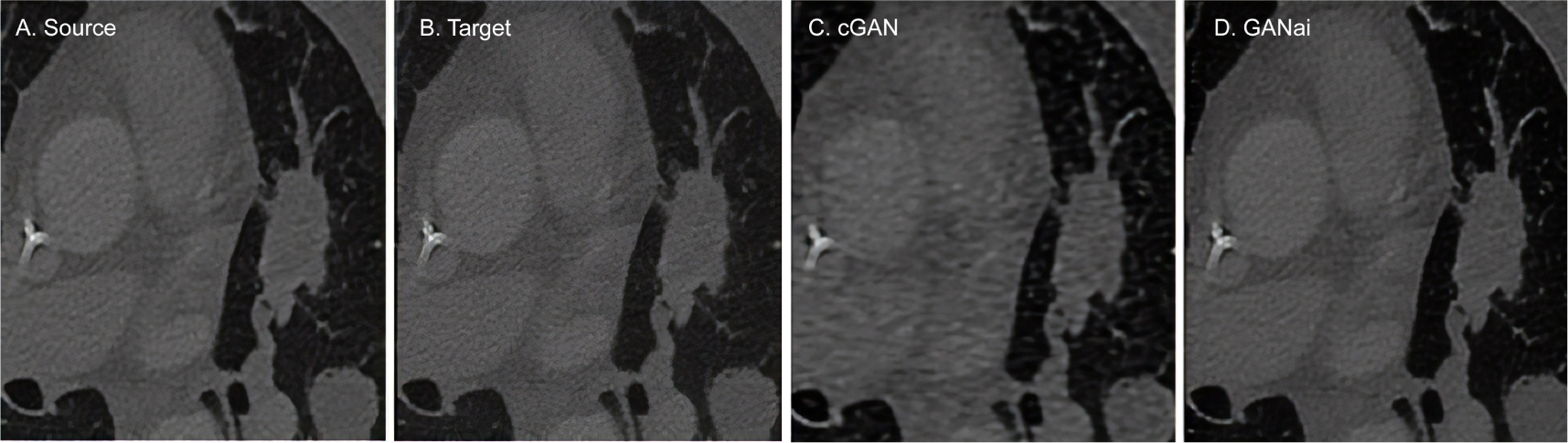
Examples of the synthesized images generated by cGAN and GANai at 100th epoch. (A) source image. (B) target image. (C) cGAN synthesized image. (D) GANai synthesized image.

## V. Discussion

### A. Training Effectiveness

The training data in GANai are separated into two subsets for the training of *G* and *D*. Our assumption is that for certain source images that are difficult to standardize, we should avoid them in the *G*-training phase. Instead, we use them to train *D*. To test the assumption, we trained a new GANai model called *GANai*_*reverse*_ with the opposite training data assignment (i.e., *G* trained with *T*_*hard*_ and *D* trained with *T*_*easy*_). Figure 10 shows that the mean absolute errors of *GANai*_*reverse*_ are significantly higher than GANai on a majority of the features, indicating that training data assignment is critical for improving GAN performance.

**Fig. 10:**
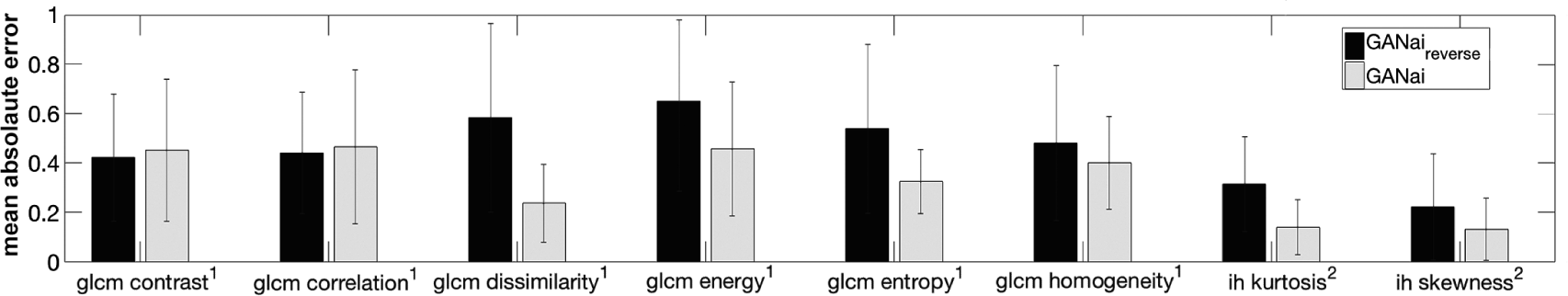
Averaged feature errors for the data effectiveness test. ^1^ Gray-level co-occurrence matrix, ^2^ Intensity Histogram. It shows that the mean absolute errors of GANai_*reverse*_ are significantly higher than GANai on a majority of the features, indicating that training data assignment is critical for improving GAN performance.

We further tested the effectiveness of the new strategies developed for improving training effect. Two modified c-GAN models were trained, one with dedicated training data, i.e., *T*_*hard*_ for *D* and *T*_*easy*_ for *G*, called *cGAN*_*SpDa*_, and the other further adopting the alternative training strategy, called *cGAN*_*SpDa+AI*_. Experimental results show that 1) *cGAN*_*SpDa*_ can effectively reduce the feature-based absolute errors of *cGAN* on a majority of the texture features, and 2) *cGAN*_*SpDa+AI*_ can further reduce the absolute errors on texture features (Figure 11). It indicates that the new training strategies developed in GANai are effective and can be adopted by generic GAN models to further improve their performance.

**Fig. 11:**
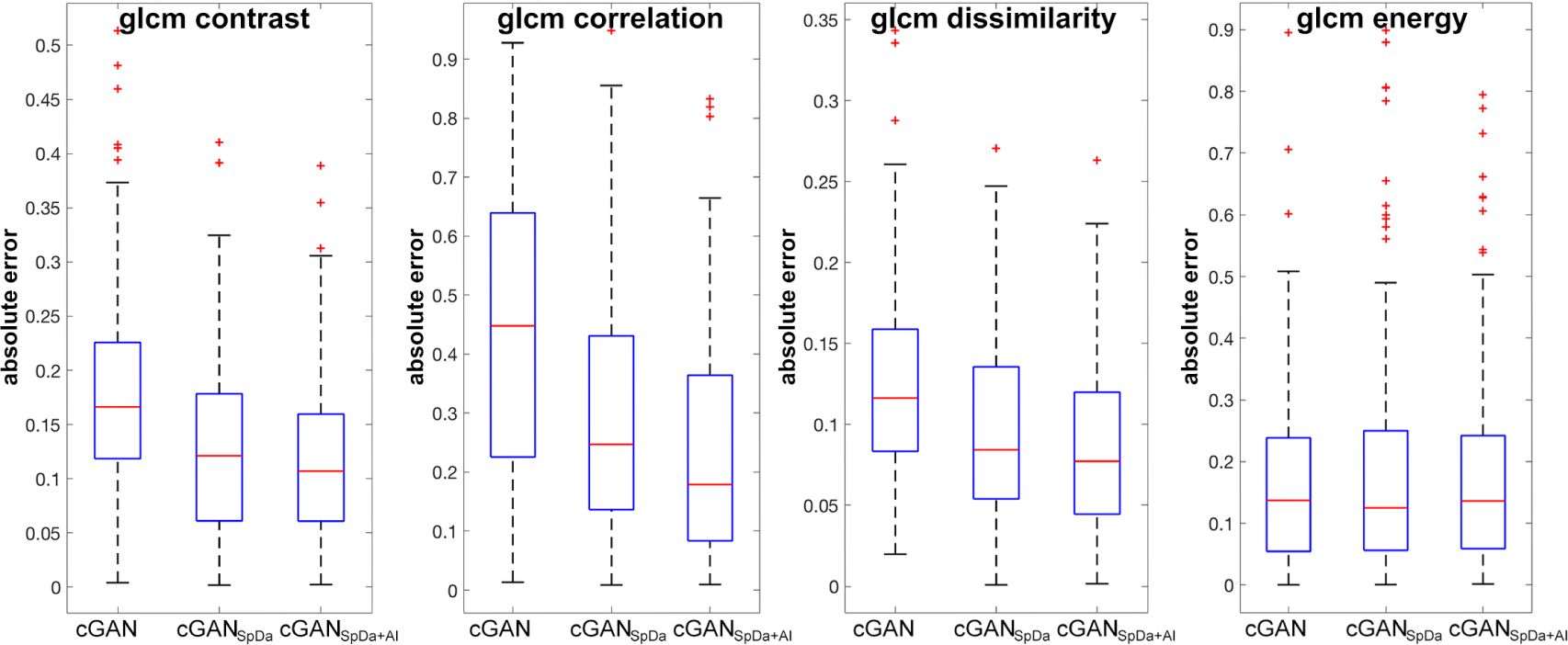
Gray-level co-occurrence matrix feature errors of different cGAN versions. *cGAN*_*SpDa*_ was trained with dedicated training data. *cGAN*_*spDa+Ai*_ was further adopting the alternative training strategy.

### B. Effectiveness of Ensemble Learning

GANai adopts the alternatively improving strategy to train *D* and *G* so that both modules can be optimized in each iteration of training. One potential problem of such full optimization is that the model could be trapped at the local minima instead of reaching the global optimization. One such example is shown in Figure S14, where a generator has been trained for more than five epochs, but it still did not result in any significant improvement. It is reasonable to believe that the model was trapped at a local minima. To address this issue, we adopt the ensemble learning approach, i.e., GANai requires a *D* to discriminate multiple *G*s and a *G* to fool multiple *D*s. Also, we rollback to the previous training phase and then retrain the model, if a satisfactory loss cannot reach in a reasonable amount of time.

**Fig. 9:**
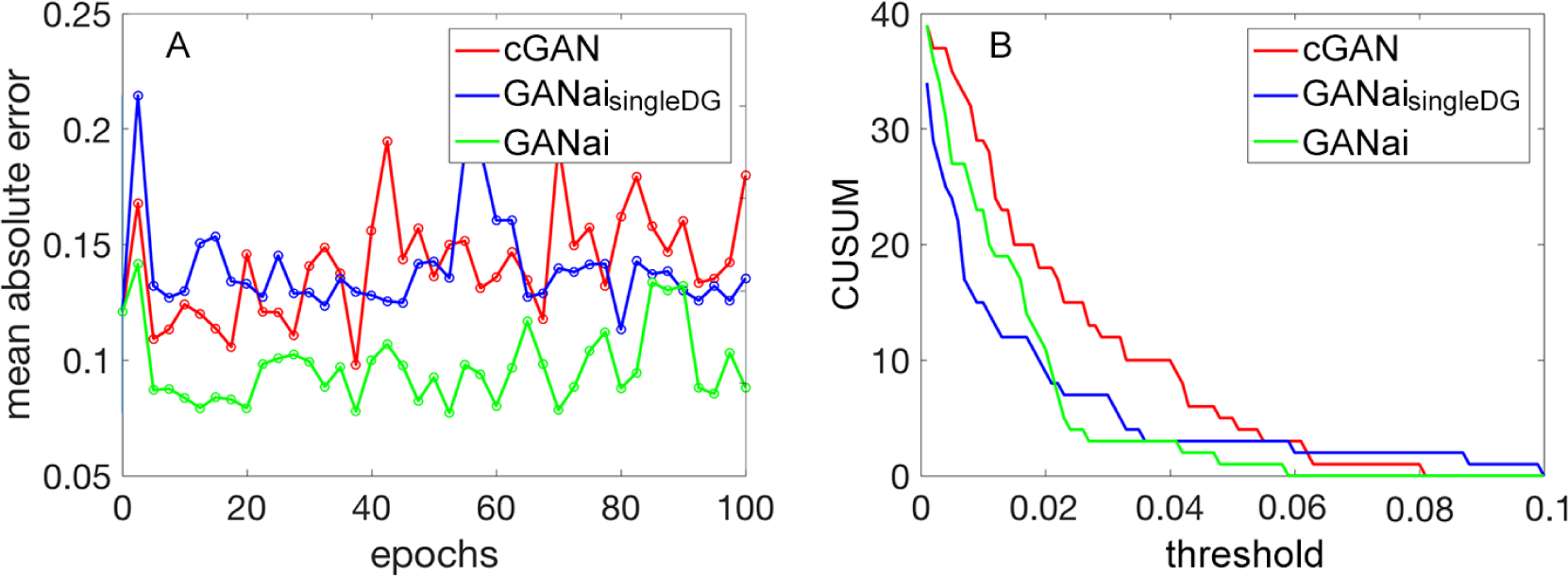
Performance evaluation on training stability. (A) the mean absolute errors of cGAN, *GANai*_*singieDG*_, and GANai on the contrast feature computed using the gray-level co-occurrence matrix. (B) the CUSUM values of cGAN, *GANai*_*singleDG*_, and GANai, where the x-axis is the threshold of CUSUM, and the y-axis is the CUSUM value. In general, GANai is the most stable model among the three.

### C. Validation Loss

The validation loss of *G* in Figure 5B is constantly lower than the training loss, which is uncommon to machine learning tasks. This is reasonable because the loss of *G* is −log*D*(*x*, *x*′) computed using the prediction result on all the fake image pairs. As shown in Figure 4, the value of *D*(*x*, *x*′) on the validation dataset is higher than that on the training dataset. After taking the minus log, the validation loss is smaller than the training loss. However, as stated in Gulrajani et al [47], the loss of GANs may not associate with model performance. Thus, the fact that the validation loss of *G* is smaller than the training loss does not necessarily indicate whether the synthesized images on the validation dataset is better than that on the training dataset. It is also why GANai uses the prediction of *D*, rather than using the loss of *G*, to control the model training phase switch.

### D. Limitations

While GANai, in general, performs better than traditional GAN models and histogram matching on texture features, its performance could be suboptimal on shape-based features. Shape-based features, such as volume, are usually determined by the physical setup of CT machines. For instance, a 1.5 mm nodule can be totally omitted in a 3 mm slice thickness scan due to partial volume [48].

## VI. Conclusion

As a popular diagnostic image modality, CT is routinely used for assessing anatomical tissue characteristics. However, CT imaging customization poses a fundamental challenge in radiomics, since non-standardized imaging protocols are commonplace. Image synthesis algorithms have been developed to integrate and standardize CT images. Among them, GAN models learn the data distribution of training data and generate synthesized images under the same distribution of the training images. However, GANs are not directly applicable to the CT image mitigation task due to the lack-of-detail problem.

We developed a novel GAN model called GANai to mitigate the differences in radiomic features of CT images. Given source images, GANai composes synthesized images by specifying a high-level goal that the image features of the synthesized images should be similar to those of the target images. GANai introduces the alternative training strategy to GAN. In each training phase, the model aims to optimize either *G* or *D* while freezing the other component. A training phase will stop if the current component is well trained or the training step exceeds an upper bound. After that, GANai switches to train the counter component. Note that just because of the adoption of the alternative training strategy, new technical improvements become applicable. For example, the inputs of the ensemble learning (multiple states of *D*s and *G*s) are the end products of every alternative training phase, and a new loss function and dedicated training data can be specified in different training phases. GANai was compared with the start-of-the-art cGAN model [22] and the patch-based histogram matching method [16]. The experimental results show that GANai is significantly better than cGAN and patch-based histogram matching on the texture and intensity histogram based radiomic features.

In conclusion, GANai is a new GAN model for CT image standardization. Its alternative training strategies are effective, easy to implement, and can be adopted by the other GAN models to further improve their performance. With GANai, CT images from multiple medical centers can be seamlessly integrated and standardized, and large-scale radiomics studies can be conducted to extract comprehensive radiomic features and to identify key tumor characteristics that drive disease transformation, progression, and drug resistance.

